# Transcriptome analysis of the necrotrophic pathogen *Alternaria brassicae* reveals a biphasic mode of pathogenesis in *Brassica juncea*

**DOI:** 10.1101/2022.09.12.507536

**Authors:** Sivasubramanian Rajarammohan

**Author notes:** Address correspondence to Sivasubramanian Rajarammohan,.

## Abstract

*Alternaria* blight or leaf spot caused by *Alternaria brassicae* has an enormous economic impact on the *Brassica* crops grown worldwide. Although the genome of *A. brassicae* has been sequenced, little is known about the genes that play a role during the infection of the host species. In this study, the transcriptome expression profile of *A. brassicae* during growth and infection was determined. Differential expression analysis revealed that 3921 genes were differentially expressed during infection. Weighted gene co-expression network analysis helped identify nine modules, which were highly correlated with growth and infection. Subsequent gene ontology (GO) enrichment analysis of the modules highlighted the involvement of biological processes such as toxin metabolism, ribosome biogenesis, polysaccharide catabolism, copper ion transport, and vesicular trafficking during infection. Additionally, 194 CAZymes and 64 potential effectors were significantly upregulated during infection. Furthermore, 17 secondary metabolite gene clusters were also differentially expressed during infection. The clusters responsible for the production of Destruxin B, Brassicicene C, and HC-toxin were significantly upregulated during infection. Collectively, these results provide an overview of the critical pathways underlying the pathogenesis of *A. brassicae* and highlight the distinct gene networks that are temporally regulated, resulting in a biphasic mode of infection. The study thus provides novel insights into the transcriptional plasticity of a necrotrophic pathogen during infection of its host. Additionally, the *in planta* expression evidence for many potential effectors provides a theoretical basis for further investigations into the effector biology of necrotrophic pathogens such as *A. brassicae*.

## INTRODUCTION

*Alternaria* blight or leaf spot is endemic to the Brassica crops grown in the tropical and sub-tropical regions worldwide. It is caused by a wide variety of pathogenic species from the ubiquitous fungal genus *Alternaria* (1). Among the *Alternaria spp*. invading *Brassicaceae, A. brassicae, A. brassicicola, A. alternata*, and *A. raphani* reportedly cause the most damage (2). *(A) brassicae* is a necrotrophic pathogen that affects its host species at all stages of growth and symptoms of infection. *A. brassicae* is one of the dominant invasive species on oleiferous Brassicas. *Brassica juncea* is widely grown in the Indian subcontinent, Australia, and parts of Europe as an oilseed crop. *A. brassicae* is a major constraint to realizing the yield potential of *(B) juncea* in the Indian subcontinent. Understanding the molecular mechanisms employed by the pathogen during the infection is essential for identifying novel targets for disease management. The pathogenicity of the related species, *A. alternata*, has been attributed to secondary metabolites or host-specific toxins (3). Although several toxins, effectors, and virulence factors of *A. brassicicola* have been characterized, the mechanism by which it colonizes the host remains unknown (4). However, the knowledge of the mechanisms/pathways underlying the pathogenesis of *A. brassicae* is limited. Studies on the molecular aspects of *A. brassicae* pathogenicity have primarily concentrated on the roles of secondary metabolite toxins such as Destruxin B (5, 6). The study of other pathogenicity factors in *A. brassicae* was precluded by the hypothesis that Destruxin B was a host-specific toxin, and detoxification of the toxin could lead to the abolition of infection (7). Recent studies confirmed that Destruxin B is not a host-specific toxin, and other proteinaceous toxins/effectors may be responsible for the pathogenicity of *A. brassicae* (8).

High-quality genomes of various *Alternaria* species are currently available with the advent of newer and cheaper sequencing technologies (13-16). While this has accelerated the study of pathogen biology and the identification of various effectors and pathogenicity factors, functional evidence for most identified genes in the form of transcriptomic data is lacking. Transcriptome analysis of *Alternaria* leaf spot infections has primarily focused on the plant responses to infection (9-12). Additionally, these transcriptome studies were performed with cDNA microarrays; hence, retrieving fungal reads/transcripts from the raw data is not feasible.

Given the enormous economic impact of *A. brassicae* worldwide and the lack of a comprehensive transcriptomic profile of *A. brassicae* during infection, we undertook the current study to 1) determine the transcriptome expression profile of *A. brassicae* during growth and infection, 2) identify and functionally annotate the differentially expressed genes (DEGs) during different infection stages, 3) identify and classify gene co-expression networks that are differentially regulated temporally, and 4) analyze the repertoire of CAZymes, secondary metabolite encoding gene clusters, and effectors that are differentially expressed *in planta* during infection.

## RESULTS

### Sequencing and Assembly

*A. brassicae* is a slow-growing pathogen compared to other phytopathogens of the *Alternaria* genus. Therefore, we collected seven-day-old mycelia from the potato dextrose agar (PDA) plates to obtain enough biomass for RNA extraction. *A. brassicae* spores maximally penetrate the host tissue at 2 days post inoculation (dpi) (17). Therefore, 2 dpi and 4 dpi were designated as the early and late time points of infection, respectively. The maximum number of droplets were inoculated on the leaf surface to obtain an optimal yield of *in planta* fungal mycelia (Fig. 1B). RNA from both plate-grown fungi (*in vitro*) and infected leaf tissue (*in planta*) were sequenced by paired-end sequencing, yielding approximately 218 million reads (Fig. 1A).

**FIG 1:**
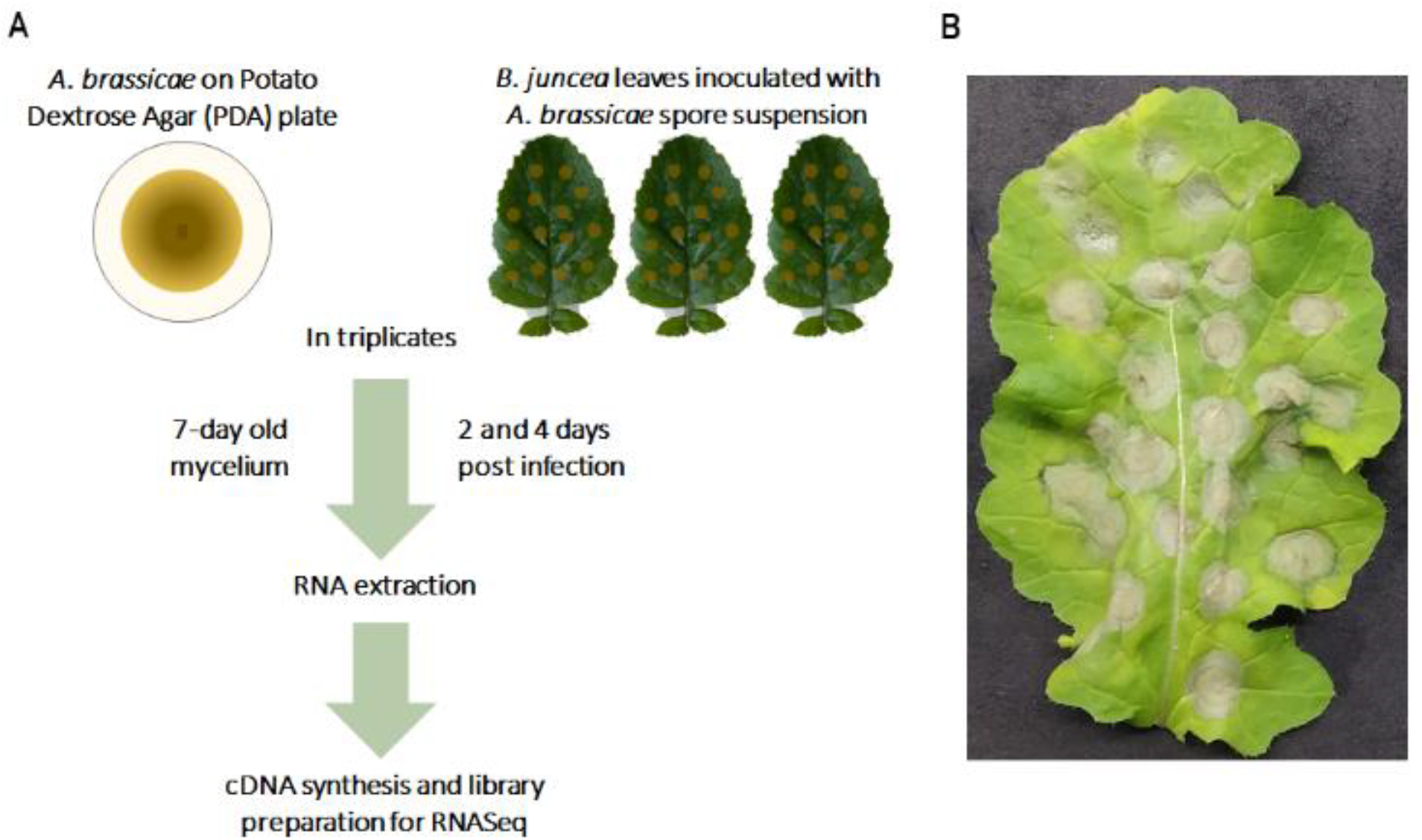
An overview of the transcriptome analysis experimental design. (A) Schematic representation of the sample collection and RNA sequencing strategy for the transcriptome analysis of *in vitro* and *in planta* samples. (B) Representative image of *B. juncea* leaf demonstrating symptoms of *A. brassicae* infection (four days post-infection; 4 dpi).

Approximately 92.8% of the reads from the *in vitro* samples mapped to the *A. brassicae* reference genome. In contrast, 0.2–1.6% of the 2-dpi *in planta* samples and 1.3–3.1% of the 4-dpi *in planta* samples mapped to the *A. brassicae* genome (Table S1). The lower percentage of mapping in *in planta* samples has also been observed in other pathogen-host interaction studies (18-21). A gene was defined to be expressed in a particular sample if > 10 reads from a sample type could be mapped to the gene. A total of 11,124 (95.9%) genes were found to have detectable expression levels in at least one of the sample types (*in vitro* or *in planta*) of which 8603 genes (77.3% of the expressed genes) were expressed in the *in planta* samples. Despite the low number of fungal reads in the *in planta* samples, they mapped to >75% of the total expressed genes of *A. brassicae*. Principal component analysis suggested that differences in gene expression could be majorly attributed to the sample type (*in vitro* vs. *in planta*, PC1: 66%) and time of sampling (PC2: 20%) (Fig. 2). Furthermore, the biological replicates of each sample type clustered together indicating high correlation within the samples of each type (*in vitro*, 2 dpi, and 4 dpi).

**FIG 2:**
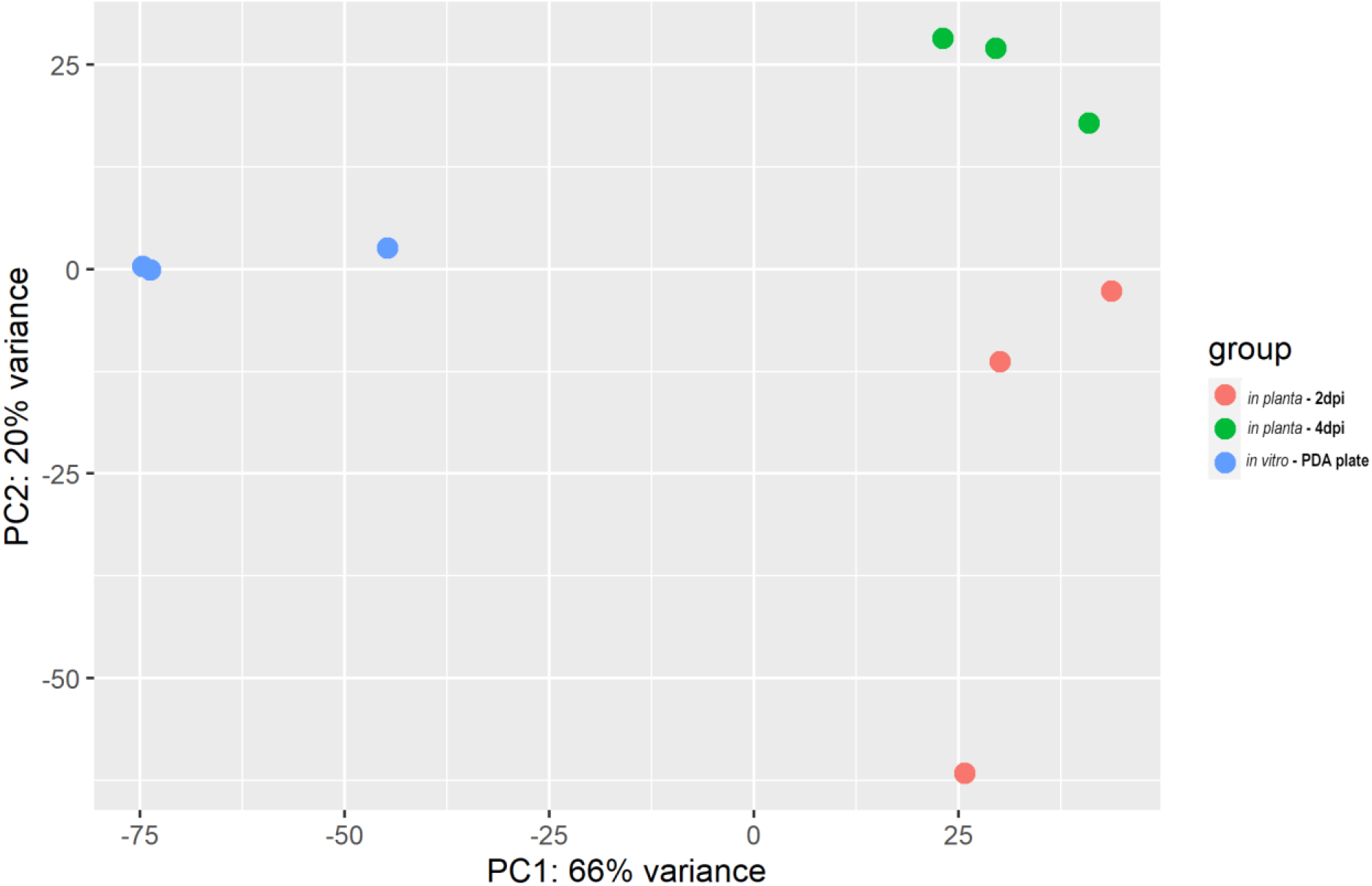
Principal component analysis. The PCA plot for transcriptome expression data depicting the clustering by sample type (*in vitro* vs. *in planta*) and time of sample collection (early - 2dpi and late – 4dpi). The PCA analysis was carried out using the DESeq2 package in R.

### Differential gene expression analysis of *A. brassicae* transcriptome

Differential gene expression analysis was performed to identify *A. brassicae* transcriptome changes during the pathogenesis of *B. juncea*. A total of 2490 and 3159 *A. brassicae* genes were significantly differentially expressed at the early (2 dpi) and late (4 dpi) stages of infection, respectively. Among these, 1167 and 1323 genes were up- and downregulated, respectively, at 2 dpi. Similarly, 1569 and 1590 genes were up- and downregulated, respectively, at 4 dpi (Fig. 3A; Table S2).

**FIG 3:**
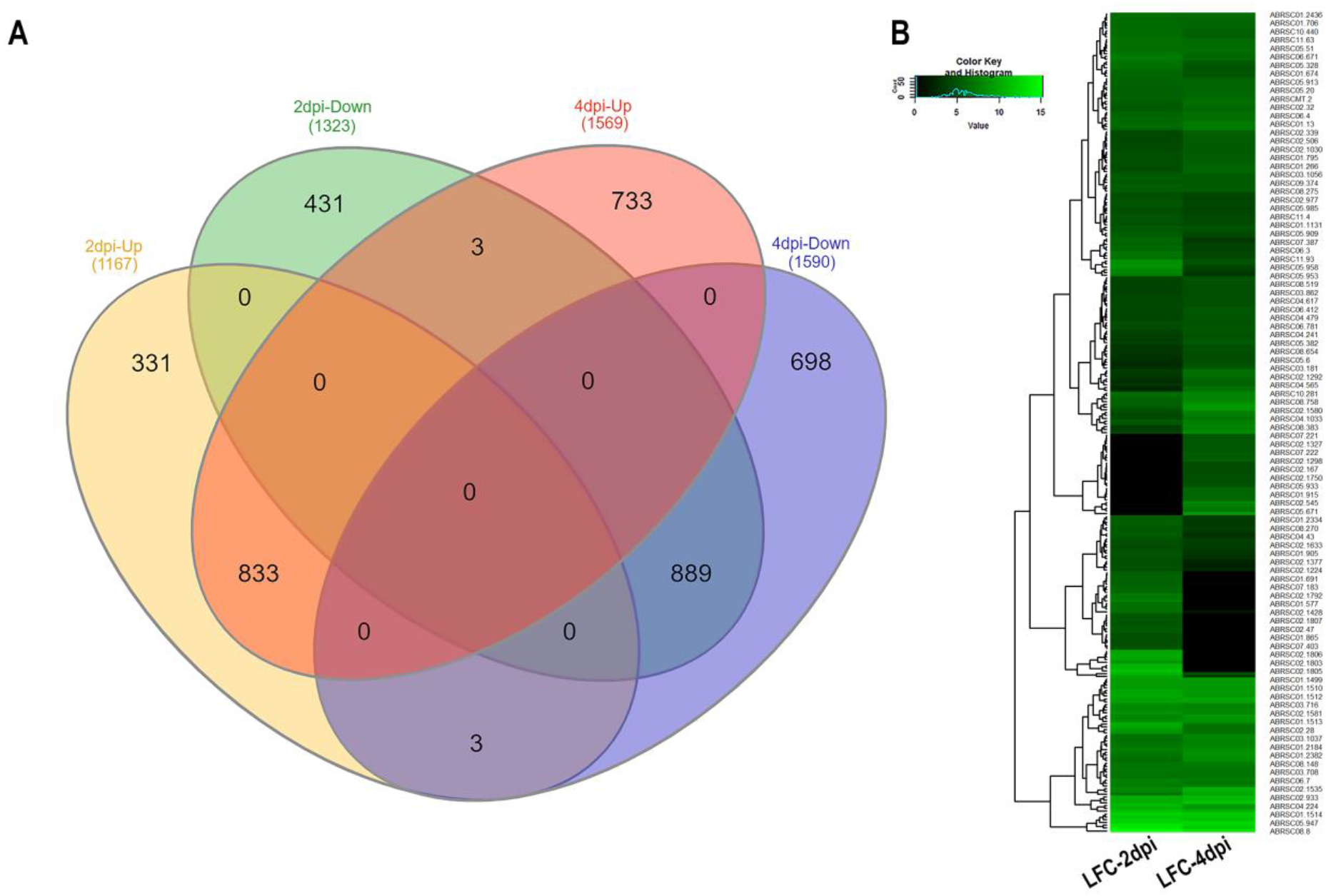
Differentially expressed genes of *A. brassicae*. (A) Venn diagram representing the number of common and unique genes between up and downregulated gene sets of early and late stages of infection. (B) Heatmap depicting the highly upregulated genes (> 25-fold; LFC > 4.6) at 2 and 4 dpi. The genes are clustered according to their expression levels.

A total of 210 and 209 genes were up-regulated by >25-fold (LFC > 4.6) at 2 and 4 dpi, respectively. Among these highly up-regulated genes, 127 were consistently upregulated by > 25-fold at 2 and 4 dpi (Fig. 3B; Table S3). Furthermore, we also identified 331 and 733 genes that were specifically upregulated at 2 dpi and 4 dpi, respectively. Three genes viz. ABRSC02.49, ABRSC07.423, and Scaffold18.17 were significantly upregulated at 2 dpi but were downregulated at 4 dpi. In contrast, three genes, ABRSC02.163, ABRSC01.1330, and ABRSC09.497, were significantly downregulated at 2 dpi but were upregulated at 4 dpi.

### Functional annotation of DEGs

Gene functional enrichment analysis was performed by assigning gene ontology (GO) terms to DEGs. DEGs were grouped according to their putative roles in biological process (BP), molecular function (MF), and cellular component (CC) as per the GO consortium. GO enrichment analysis of DEGs at 2 and 4 dpi provided insights into the biological pathways that were temporally regulated during pathogenesis.

At the early infection stage, GO BP terms such as ribosome biogenesis, ribosome assembly, carbohydrate metabolism, pectin catabolism, toxin biosynthesis, and RNA processing were enriched in the differentially up-regulated genes. GO BP terms such as negative regulation of chromosome organization, negative regulation of DNA-templated transcription, cellular protein modification, autophagy, and amino acid transport were enriched in the downregulated genes at the early infection stage (Fig. 4; Table S4).

**FIG 4:**
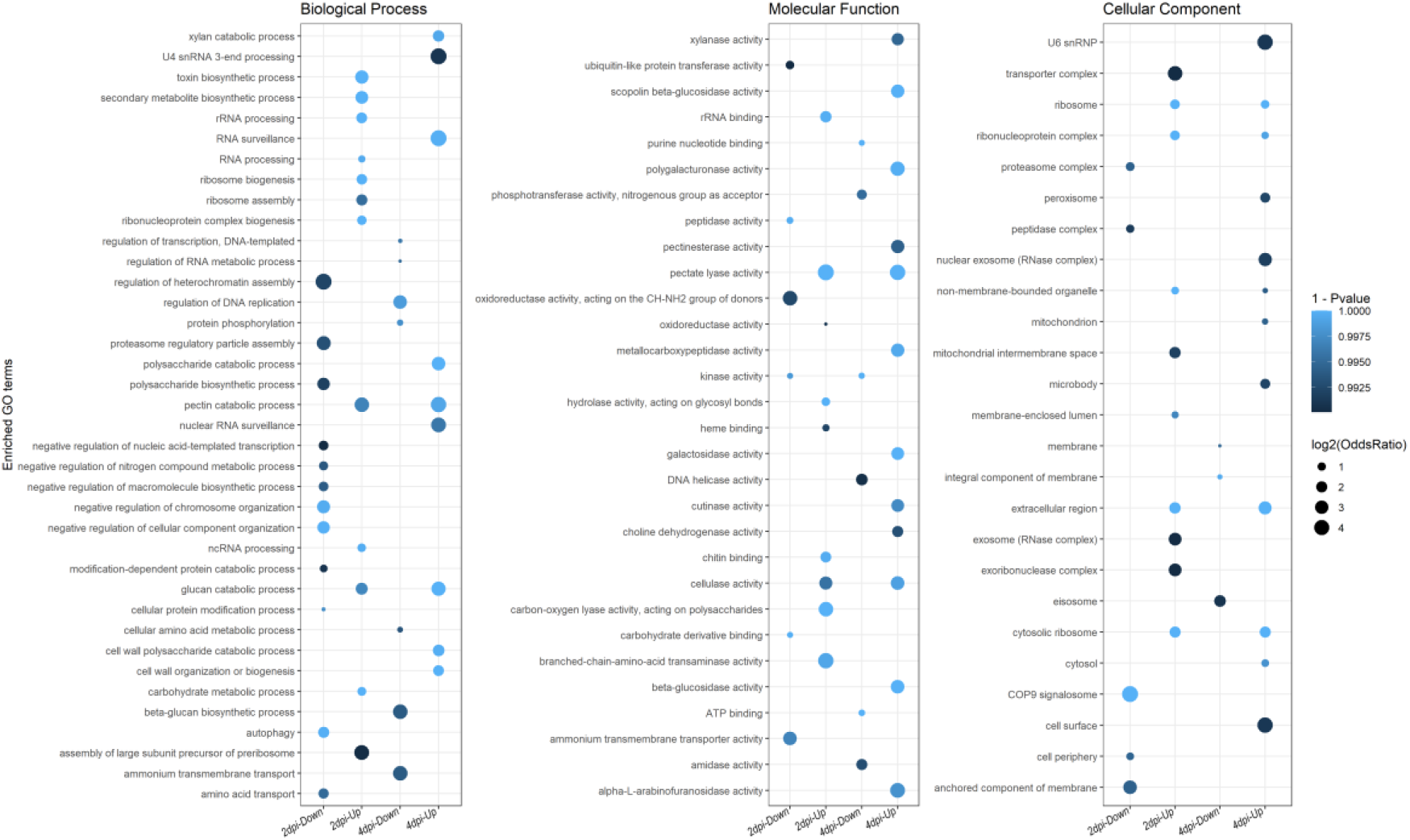
GO enrichment analysis of DEGs. Dotplots depicting the GO enrichment of DEGs under biological process, molecular function, and cellular component categories. The color of the dots represents the p-value of the hypergeometric test for GO enrichment, and the size of the dot represents the odds ratio of the GO term with respect to that in the background gene set.

Similarly, at the late infection stage, GO BP terms such as cell wall polysaccharide catabolism, cell wall organization, xylan/pectin catabolism, and RNA surveillance were enriched in the differentially upregulated genes. GO BP terms such as regulation of DNA replication, protein phosphorylation, regulation of transcription, amino acid metabolism, and ammonium transport were enriched in the differentially down-regulated genes at the late infection stage (Fig. 4). Additionally, the molecular functions associated with the above processed were also similarly enriched in the respective gene sets (Fig. 4; Table S5).

### Weighted gene co-expression network analysis (WGCNA) reveals a biphasic mode of infection

To identify functional pathways involved in the pathogenesis of *A. brassicae*, highly co-expressed gene modules were inferred from the DEGs using WGCNA. A total of 3921 DEGs were used to construct the co-expression network. In WGCNA, modules are defined as clusters of highly interconnected genes possibly having a functional relationship. Using a softpower threshold of 18 (R^2^>0.9) and a minimum module size of 50, 12 modules with module sizes ranging from 79–1680 genes were identified (Fig. S1, Fig. S2; Table S6). The gene modules were correlated to the sample traits, i.e., growth on the PDA plate, early infection (2 dpi), and late infection (4 dpi). We observed that 9 out of 12 modules were strongly correlated to at least one of the traits (Table S7). The black (M2) and orange (M1) were positively correlated to the growth on PDA plates, whereas the salmon (M5) module was negatively correlated to the growth on PDA. The darkgreen (M7), lightcyan (M6), and brown (M8) modules were positively correlated to the early infection stage, whereas the cyan (M9) module was positively correlated to the late infection stage. The darkred (M3) module was positively correlated to growth on PDA but was negatively correlated to the late infection stage. Similarly, the magenta (M4) module was positively correlated to the growth on the PDA but was negatively correlated to the early infection stage.

GO enrichment analysis of the genes in the M1, M2, M3, and M4 modules revealed a significant overrepresentation of GO BP terms such as cell morphogenesis, lipid catabolism, autophagy, regulation of DNA replication, tetrahydrofolate interconversion, asparagine biosynthesis, sporulation, proteolysis, endosome organization, cytosolic transport, regulation of RNA metabolism, transcription, and cell communication (Fig. 5A, Fig. S3–5; Table S8). The overrepresentation of these pathway genes indicates that the transcriptional machinery is geared toward cell division and primary metabolism during the growth on the PDA.

**FIG 5:**
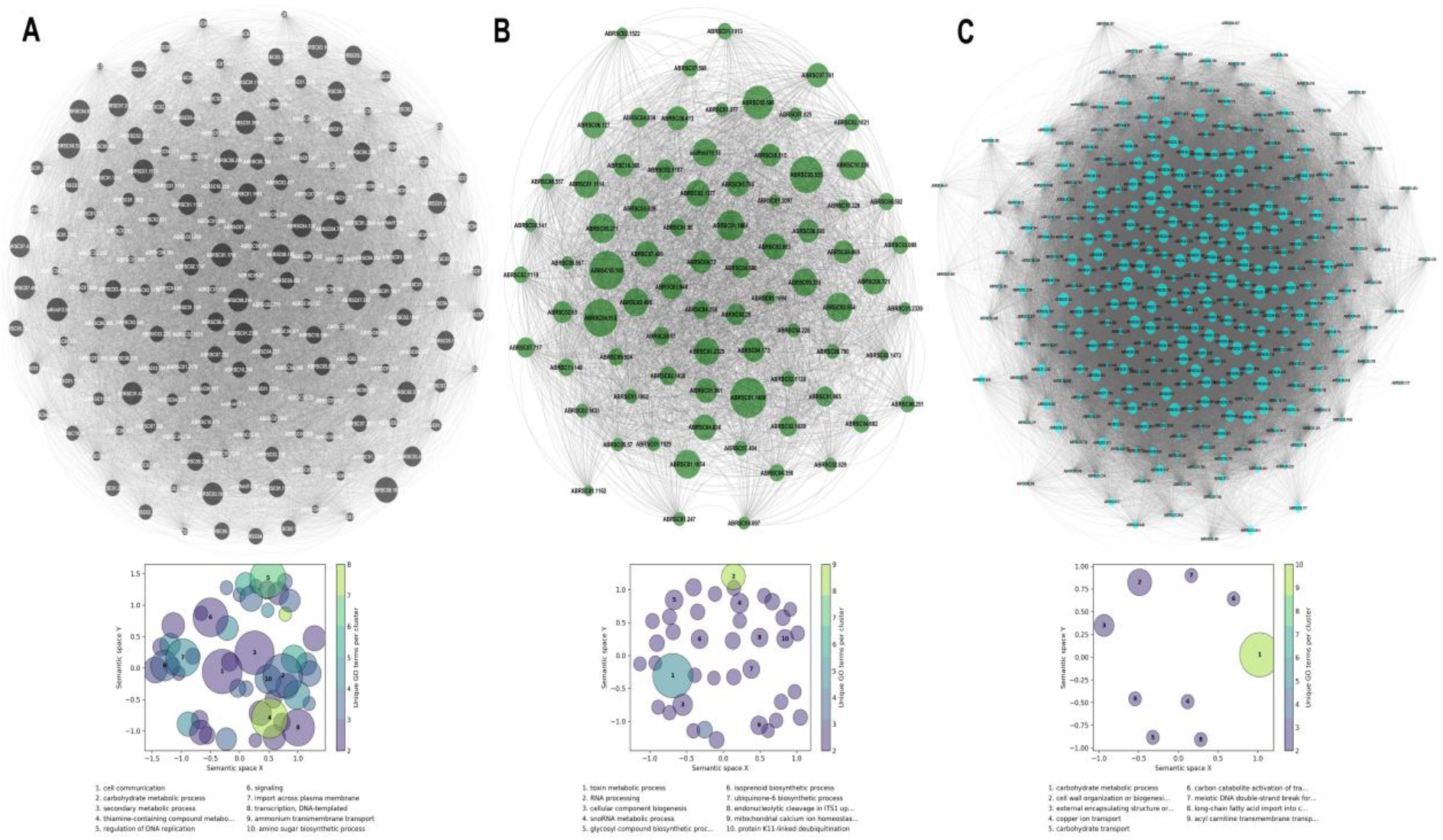
WGCNA and GO enrichment analysis of DEGs. (A) Weighted gene co-expression network of the genes in the black module and GO terms enriched in the modules correlated with growth on PDA. (B) Weighted gene co-expression network of the genes in the darkgreen module and GO terms enriched in the modules correlated with the early infection stage (2 dpi). (C) Weighted gene co-expression network of the genes in the cyan module and GO terms enriched in the modules correlated with the late infection stage (4 dpi).

In contrast, in the M6, M7, and M8 modules, GO BP terms such as glycosyl compound biosynthesis, toxin biosynthesis process, secondary metabolite biosynthesis, RNA processing, polyadenylation, and methylation were overrepresented (Fig. 5B, Fig. S6–7; Table S8). Furthermore, the M9 module contained a significant overrepresentation of GO BP terms such as carbohydrate metabolic process, xylan/pectin catabolic process, vesicular trafficking, and cell wall organization (Fig. 5C; Table S8).

The network analysis followed by the GO enrichment analysis exhibits a clear demarcation in the pathways enriched at the three different stages studied. The growth phase on PDA was enriched for pathways that governed cell growth, division, and primary metabolism. The infection process *in planta* was also bifurcated into two distinct phases wherein the early stage (2 dpi) was enriched for pathways involved in RNA processing, ribosomal translation, and toxin metabolism. In contrast, the pathways at the late stage were predominantly enriched for degradation of the host cell-wall components and vesicular trafficking.

### Candidate genes involved in the pathogenesis of *A. brassicae*

#### CAZymes

Most necrotrophic pathogens secrete various carbohydrate-active enzymes (CAZymes) to degrade and break down the plant cell wall. We cataloged the CAZymes that were significantly upregulated during the pathogenesis of *A. brassicae*. We identified 118 and 187 CAZymes to be significantly upregulated at 2 and 4 dpi, respectively. However, only 86 and 136 CAZymes significantly upregulated at 2 and 4 dpi, respectively, were secreted outside the cell. A total of 194 unique CAZymes were found to be significantly upregulated during the course of infection. The 194 CAZymes included 100 glycoside hydrolases (GH), 11 glycosyl transferases (GT), 21 carbohydrate esterases (CE), 15 carbohydrate-binding modules, 30 auxiliary activity enzymes, and 17 polysaccharide lyases. Polysaccharide lyases (PLs) cleave uronic acid-containing polysaccharides via a β-elimination mechanism and are responsible for the degradation of pectin, one of the most abundant components of the plant cell wall. We identified 14 secretory PLs that were highly upregulated at both 2 and 4 dpi (Table S9). However, we observed that polygalacturonases (GH28 family) that are responsible for hydrolyzing the α-1,4 glycosidic bonds in pectin are exclusively upregulated during the late stage of infection (4 dpi) (Table S9).

#### Effectors

Effectors are small secretory proteins with distinct physicochemical properties that play an important role in establishing pathogenesis. Computational prediction of effectors in the *A. brassicae* genome revealed 198 putative candidates (22). We further examined the expression status of these 198 putative effectors during infection. We observed that 116 of the 198 effectors were significantly upregulated during infection (Table S10; Fig. S8). Consistent expression across both the time points was observed for 64 of the 116 effectors expressed during infection. A total of 70 and 110 of the 116 effectors were significantly upregulated at 2 and 4 dpi, respectively. Additionally, we also observed that 45 of the 116 effectors were also characterized as CAZymes, thus indicating their pivotal role in the pathogenesis of *A. brassicae*. The AA9 family or the copper-dependent lytic polysaccharide monooxygenases (LPMOs) were reportedly expanded in the *Alternaria* genus (22). We identified eight of the 11 previously reported secreted LPMOs in *A. brassicae* to be significantly upregulated at 4 dpi. However, only two of the LPMOs were significantly upregulated at 2 dpi. Other known effectors such as the conserved fungal extracellular membrane (CFEM) domain-containing proteins, Nudix effectors, necrosis and ethylene-inducing proteins (NLPs) were found to be significantly upregulated during 2 and 4 dpi (Table S10). We observed that both the previously characterized AbrNLPs (23) were significantly upregulated at 2 and 4 dpi.

#### Toxins/Secondary metabolites

Secondary metabolites or non-proteinaceous toxins are known pathogenicity factors of many phytopathogens, including *Alternaria alternata*. Toxins are encoded by biosynthetic gene clusters that are usually co-regulated. *A. brassicae* genome was predicted to contain 34 secondary metabolite (SM) gene clusters (22). We evaluated the expression profile of the 34 SM clusters during growth and infection. A total of 17 SM gene clusters were differentially expressed during infection, among which 10 SM gene clusters were significantly upregulated and seven SM clusters were significantly downregulated. The siderophore (dimethylcoprogen) and melanin gene clusters were significantly downregulated during infection (Fig. 6, Table S11). In contrast, the Destruxin B, Brassicicene C, and HC-toxin gene clusters were significantly upregulated during infection (Fig. 6; Table S11). Additionally, several other SM gene clusters involved in the synthesis of unknown metabolites/toxins were also significantly upregulated during infection.

**FIG 6:**
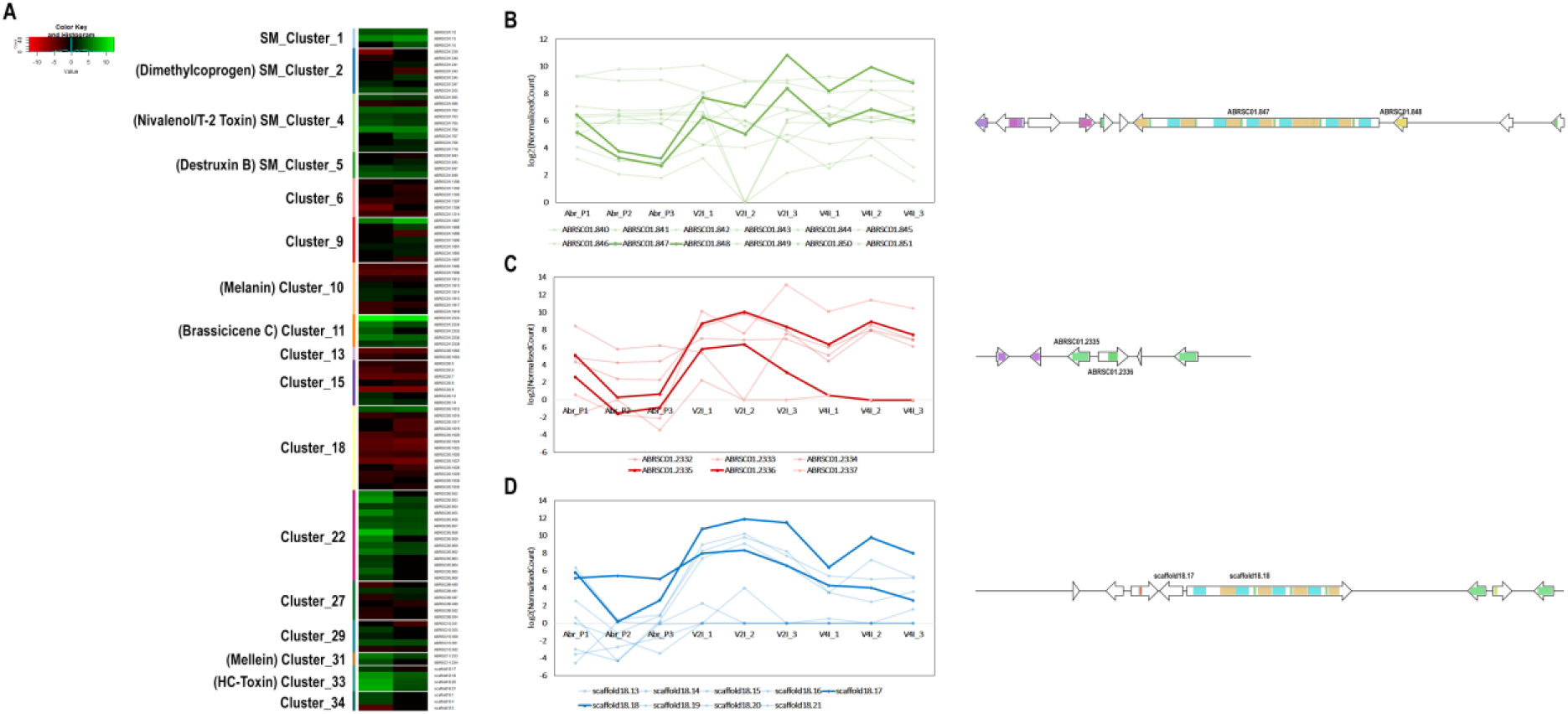
Expression profiles of secondary metabolite gene clusters in *A. brassicae*. (A) Heatmap indicating the expression profiles of significantly up- or downregulated gene clusters. Normalized expression values of the SM cluster genes encoding (B) Destruxin B, (C) Brassicicene C, and (D) HC-toxin in the nine samples. The lines denoting the core biosynthetic genes in (B), (C), and (D) are darkened.

## DISCUSSION

In the present study, we investigated the alterations in the transcriptome profiles of *A. brassicae* growth on an artificial medium and during the infection of *B. juncea*, its natural host. Transcriptome studies on host-pathogen interactions are limited by the challenges in obtaining an adequate representation of pathogen sequences for further analyses (18-21). Therefore, we employed an inoculation method to maximize the enrichment of fungal transcripts in the infected leaf samples. Although our approach yielded fewer reads mapping to the reference genome, we obtained >75% coverage (8603 of the 11,124 genes) of the transcripts in the *in planta* samples. In total, we identified 2490 and 3159 *A. brassicae* DEGs at the early and late stages of infection. Functional annotation of the DEGs followed by GO enrichment analysis helped identify transcriptionally active key pathways/processes during different stages of infection.

The gene co-expression analysis and subsequent GO enrichment analysis have led to the identification of the infection program modulation across the two infection stages. During the course of *A. brassicae* infection, penetration through the stomata is initiated as early as 6 hours post inoculation, however maximum penetration sites are observed at 2 dpi (17). At 2 dpi, water-soaked lesions start to appear at the macroscopic level, whereas, by 4 dpi clear necrotic lesions are visible on the leaf surface (17). We found distinct changes in the sets of biological processes activated during growth and the two infection stages (Fig. 7). The colonization of *A. brassicae* caused marked changes in the transcriptional activity at 2 and 4 dpi. This is further supported by the observation that toxin metabolic processes, RNA processing, and ribosome biogenesis were enriched GO terms at 2dpi (Fig. 7).

**FIG 7:**
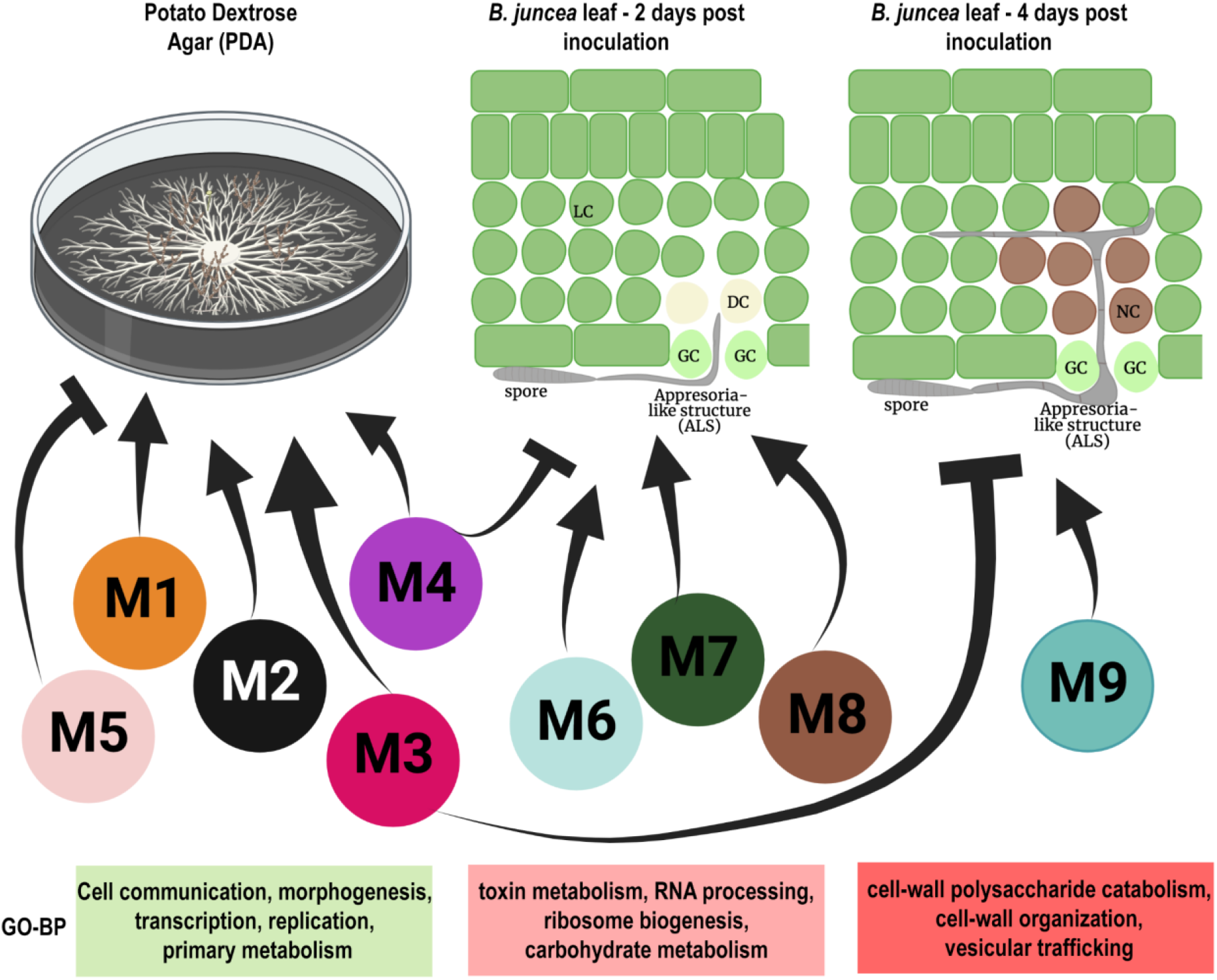
Molecular mechanisms underlying the pathogenesis of *A. brassicae* in *B. juncea*. The different WGCNA modules positively/negatively correlated with the three stages of growth and infection are shown. GC – Guard cells, DC – Dying cells, NC – Necrotic cells, and LC-Live cells. Module colors are represented per their assignment in WGCNA. Representative GO Biological Process terms of the modules are also listed.

The secondary metabolites and toxins such as Destruxin B and Brassicicene C are known to modulate plant physiology to facilitate penetration of the host and subsequently modulate the immune responses to the pathogen (6, 24). In parallel, RNA processing and ribosome biogenesis processes are required for the production of proteinaceous effectors and CAZymes that target host proteins and carbohydrates to cause necrosis and establish infection. At 4 dpi, the major GO terms that were enriched belonged to carbohydrate metabolic process, cell-wall polysaccharide catabolism, and copper ion transport (Fig. 7). This indicates the temporal shift in the infection process from facilitation of colonization to the establishment of infection by cellular degradation of the host cells. The gene co-expression networks, thus, highlighted the temporal processes unique to each infection stage. This biphasic infection process is in line with the hypothesis that most necrotrophs do not immediately cause cell death upon interaction with the host. and there may exist a short biotrophic phase before the onset of necrosis (25).

We observed that the pathogenicity factors in *A. brassicae* mostly consist of secreted CAZymes, toxins, and effectors. This is in contrast to *Alternaria alternata* pathovars, which use only specialized secondary metabolites or toxins as pathogenicity determinants (26-28). These results suggest that the necrotrophic lifestyle within the *Alternaria* genus itself has evolved through different genetic mechanisms. The plant cell wall is the primary barrier encountered by microbial pathogens, and the complexity of the plant cell wall polysaccharides determines basal resistance to most invading microorganisms. CAZymes comprise several classes of enzymes that metabolize different carbohydrate moieties (29). They are often classified as cell-wall-degrading enzymes (CWDEs) in the context of phytopathogens since they process or hydrolyze plant cell-wall polysaccharides to facilitate infection and acquire nutrients from the hydrolyzed polysaccharides (30). The diversity of the CAZymes secreted by the pathogen, in turn, affects its host range. We observed that 38% of the total CAZymes in the *A. brassicae* genome were significantly upregulated during infection. A markedly higher number of secreted CAZymes (Table S10) that are also classified as effectors point to their role as one of the main determinants of pathogenicity in *A. brassicae*. The PL and CE families were the most active enzyme families during the infection process, wherein 81% and 64% of the total PLs and CEs were expressed. Pectin is a major component in the cell wall of dicot plants and the middle lamellae that help bind cells together (31). The digestion of pectin causes tissue collapse, tissue necrosis, and subsequent release of oligopectin compounds that may trigger plant defense (32). Pectin-digesting enzymes such as pectate lyases (PL) and pectinesterases that digest pectin are, therefore, considered crucial pathogenicity factors. Loss-of-function mutations of PLs in many phytopathogens have contrasting effects on pathogenicity. In the closely related *A. brassicicola*, loss of function of two pectate lyases reportedly has opposing effects on the pathogenicity of the strain (33, 34). This indicates that functional redundancy exists in the case of some PLs, as observed in the case of other phytopathogens such as *Cochliobolus carbonum* (35). The PL family in *A. brassicae* consists of 21 members, among which 17 were significantly upregulated during infection. This indicates that the PLs in *A. brassicae* may be acting in coordination or may be functionally redundant. A systematic study of loss-of-function mutations of these PLs would be required to delineate their individual functions and their redundancy.

Secreted effectors are critical players in mediating pathogenesis. Putative effector candidates were identified in *A. brassicae* through computational approaches (22). We determined the expression profiles of the putative effector candidates to evaluate their role during the infection process. Approximately 59% of the putative effectors were expressed at least at one of the time points. This indicates that proteinaceous effectors are vital during the pathogenesis of *A. brassicae*. This is in contrast to the earlier view that secondary metabolites or toxins are major players of pathogenicity in *A. brassicae* (5-7). Although a large proportion of the *in planta*-expressed effectors do not have any known function or domain, we identified a few effectors that have been characterized previously in other phytopathogens. However, apart from AbrNLPs, no other effector from *A. brassicae* has been characterized. This study, therefore, provides *in planta* expression evidence for a majority of the putative effector candidates and lays the foundation for further functional characterization of *A. brassicae* effectors.

Pathogens belonging to the *Alternaria* and *Cochliobolus* genera are well-known producers of secondary metabolites and peptide toxins that enable them to infect their hosts. Our results reveal that some of the SM biosynthetic clusters in *A. brassicae* are transcriptionally active during infection of *B. juncea*. Some SM clusters code for known secondary metabolites, whereas most transcriptionally active clusters code for unknown or unidentified secondary metabolites. Destruxin B, a cyclic depsipeptide and a key pathogenicity factor of *A. brassicae*, is reportedly a host-specific toxin of *A. brassicae* (5). Destruxin B has not been reported to be produced by any other *Alternaria* species. Our results suggest that the core biosynthetic genes of the Destruxin B cluster, i.e., *DtxS1* and *DtxS3*, are significantly upregulated during infection. Destruxin B reportedly contributes to the aggressiveness of the pathogen but does not act as a host-specific toxin (8). The SM cluster coding for Brassicicene C was significantly upregulated during infection. Brassicicene C belongs to the Fusicoccanes family, which are diterpenoids with varying effects on plant physiology, including the opening of stomata (24). Therefore, Destruxin B and Brassicicene C may aid in the pathogenesis by modulating the plant physiology but may not be directly involved in the pathogenesis. HC-toxin is the major virulence determinant of the *Cochliobolus carbonum*, and resistance against *C. carbonum* is conferred by a carbonyl reductase gene that detoxifies HC-toxin (36). Two species of the *Alternaria* genus, viz. *A. jesenskae* and *A. brassicae*, are the only other fungal species that contain the SM cluster that produces this toxin (22, 37). The HC-toxin cluster is also one of the highly upregulated gene clusters during the infection of *B. juncea*. The presence of the HC-toxin cluster in *A. brassicae* may also be one of the factors contributing to its wider host range compared to that in the other pathogens in the *Alternaria* genus.

In conclusion, the transcriptome of *A. brassicae* infection has revealed the mechanisms by which the pathogen rewires the cellular processes during growth and infection and also highlights the transcriptional plasticity of the pathogen. In this regard, the analysis also highlights the biphasic mode of infection during which a temporal regulation of distinct gene networks exists to facilitate and establish pathogenesis. Additionally, the study has provided *in planta* expression evidence for many potential effector candidates and will thus facilitate further research into the effector biology of necrotrophic pathogens such as *A. brassicae*. Consequently, this will enable the development of novel strategies for integrated disease management in crop plants.

## MATERIAL AND METHODS

### Plant material and fungal strain

The fungal strain *A. brassicae* J3 (38) was used for the infection assays in the expression analysis. The strain was grown on potato dextrose agar (PDA, pH adjusted to 7.0 using 1 N NaOH) plates at 22°C for 15 days under a 12-h light/dark cycle. Similarly, *B. juncea* var. Varuna was grown at 25°C with a photoperiod of 10 h Light/14 h dark and used for the infection assays.

### Disease inoculation assays

Briefly, 15-day-old PDA plates were used to prepare spore suspension for drop inoculations. Multiple 15 μl droplets of the spore suspension (concentration of 3–5 × 10^3^ conidia/ml) were placed on the leaves of 5–6-week-old *B. juncea* plants to maximally cover the surface area. The leaves were then maintained at > 90% relative humidity to enable infection. Leaves were collected two and four days’ post inoculation (2 and 4 dpi). 15-day-old *A. brassicae* mycelia growing on PDA was also harvested (*in vitro* sample).

### RNA extraction, library preparation, and sequencing

Total RNA was extracted from 2–3 leaves collected from three individual plants in each experiment using the MagMAX™-96 Total RNA Isolation Kit according to the manufacturer’s recommendation (Invitrogen, Thermo Fisher Scientific, Waltham, USA). Total RNA (500 ng) was used to enrich the mRNA using NEBNext Poly (A) mRNA magnetic isolation module (New England Biolabs, Ipswich, USA) by following the manufacturer’s protocol. Libraries were prepared from the enriched mRNAs using the NEBNext® Ultra™ II RNA Library Prep Kit for Illumina (New England Biolabs, Ipswich, USA). The library concentration was determined in a Qubit3 Fluorometer using the Qubit dsDNA HS (High Sensitivity) Assay Kit (Thermo Fisher Scientific, Waltham, USA). The library quality was assessed using Agilent D5000 ScreenTape System in a 4150 TapeStation System. All the libraries were sequenced on an Illumina Novaseq platform (Clevergene Pvt. Ltd, Bengaluru, India).

### Quality control and read mapping to the reference genome

The RNA-Seq reads from nine libraries (three biological replicates per condition) were processed to remove adaptor sequences, empty reads, and low-quality sequences with a Phred score < 30 and reads < 36 bp. The processed reads were stored in FASTQ format. The reference genome and the corresponding annotations for *A. brassicae* and *B. juncea* were obtained from previous reports (16, 39). Indices for each reference genome were built, and quality filter-passed reads were mapped to the reference genomes using STAR aligner (40). Raw read counts mapped to each gene from the STAR-generated alignments were obtained using the featureCounts (41) command of the Subread package (42).

### Differential gene expression analysis

Differential gene expression (DGE) analysis was conducted using the DESeq2 package (43). The raw count data obtained from the featureCounts were used as the input for DESeq2. Principal component analysis (PCA) was conducted to determine the relatedness of the biological replicates. A false discovery rate (FDR) cutoff of 0.05 was applied to account for multiple testing corrections. Genes with a ≥ 2-fold (absolute value of log_2_ fold change |LFC| ≥ 1) change in expression level and an FDR-adjusted p-value < 0.05 were considered “differentially expressed”.

### Functional classification and enrichment analyses of DEGs

Gene ontology (GO) enrichment analysis was performed using the GOstats package (44). GO terms assigned to the total genes list of *A. brassicae* were used as the background list for enrichment analysis. The hypergeometric test implemented in GOstats was used to identify significantly enriched GO categories. A GO category was considered significantly enriched only when the FDR-corrected p-value for that category was < 0.05.

### Weighted gene co-expression network analysis of DEGs

Gene co-expression networks were constructed using the WGCNA package (45) in R. The vst transformed expression data for the DEGs were used for co-expression analysis. A soft thresholding power of 18 was determined using the pickSoftThreshold function based on the scale-free topology model fit (r2 > 0.9). The blockwiseModules function was used to obtain co-expression modules, and closely related modules were merged at a cutHeight = 0.20. Pearson’s correlation coefficient was calculated to correlate the modules with the sample traits. GOstats package was used for GO enrichment of modules. Cytoscape v3.8.2 (46, 47) was used to visualize the co-expression networks.

## DATA AVAILABILITY

The Transcriptome dataset of all the samples has been deposited in the NCBI BioProject database (BioProject number: PRJNA860050).

## SUPPLEMENTAL MATERIALS

Supplemental Tables: Supplemental Tables S1–11.

Supplemental Figures: Fig. S1–8.

## ACKNOWLEDGMENTS

This research was supported by grants from the Department of Science and Technology through the DST-INSPIRE Faculty program to SR and a Core Research Grant from the Science and Engineering Research Board (SERB Grant No. CRG/2020/001731).

## List of supplemental materials

Table S1: Read mapping statistics (reads were mapped to *A. brassicae* J3 genome)

Table S2: List of DEGs at 2 and 4 dpi of *A. brassicae* infection

Table S3: Top upregulated genes *in planta* (LFC > 4.6; Fold change > 25-fold)

Table S4: GO Biological Process terms enrichment analysis of *A. brassicae* DEGs 2 and 4 dpi

Table S5: GO Molecular Function terms enrichment analysis of *A. brassicae* DEGs 2 and 4 dpi

Table S6: WGCNA modules and their respective gene counts

Table S7: WGCNA module correlation with sample traits (plate-grown, 2 dpi, 4 dpi)

Table S8: GO enrichment analysis of WGCNA modules strongly correlated with sample traits

Table S9: Gene expression profiles of CAZymes at 2 and 4 dpi of *A. brassicae*

Table S10: Gene expression profiles of effectors at 2 and 4 dpi of *A. brassicae*

Figure S1: Softpower threshold estimation in WGCNA

Figure S2: DEGs and their module membership before and after DynamicTree Cut

Figure S3: Gene co-expression network of genes in the darkred module

Figure S4: Gene co-expression network of genes in the magenta module

Figure S5: Gene co-expression network of genes in the salmon module

Figure S6: Gene co-expression network of genes in the lightcyan module

Figure S7: Gene co-expression network of genes in the brown module

Figure S8: Gene expression profile of effectors of *A. brassicae*. (A) Heatmap depicting the log_2_ FoldChange values of 116 effectors at 2 and 4 dpi. (B) Venn diagram representing the overlap between the effectors significantly upregulated at 2 and 4 dpi.

## Notes

### Competing Interest Statement

The authors have declared no competing interest.

