## Supplementary Figures for "Transcriptome analysis of the necrotrophic pathogen *Alternaria brassicae* reveals a biphasic mode of pathogenesis in *Brassica juncea*"

Supplemental Figures S1-8.

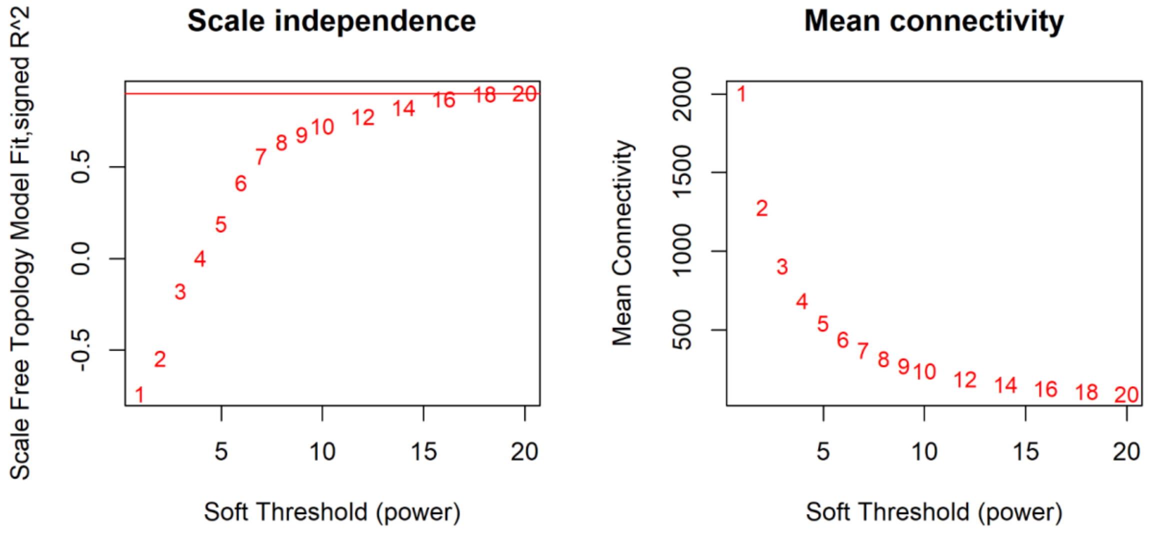


Figure S1: Softpower threshold estimation in WGCNA

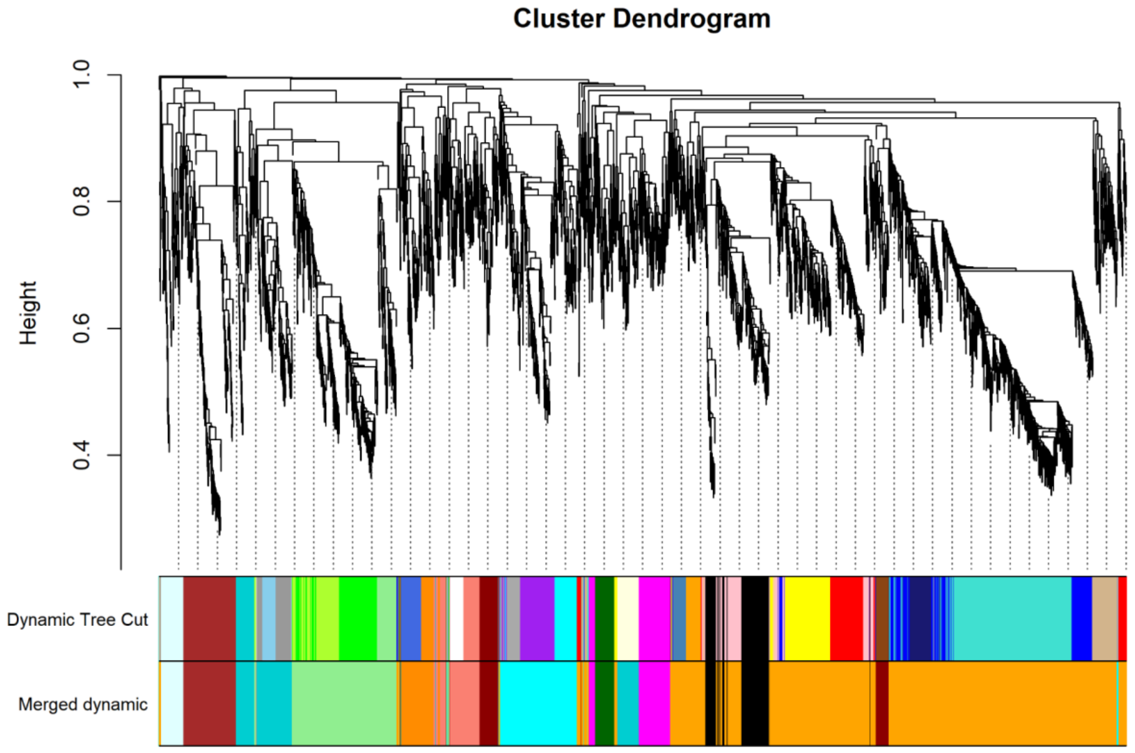


Figure S2: DEGs and their module membership before and after DynamicTree Cut

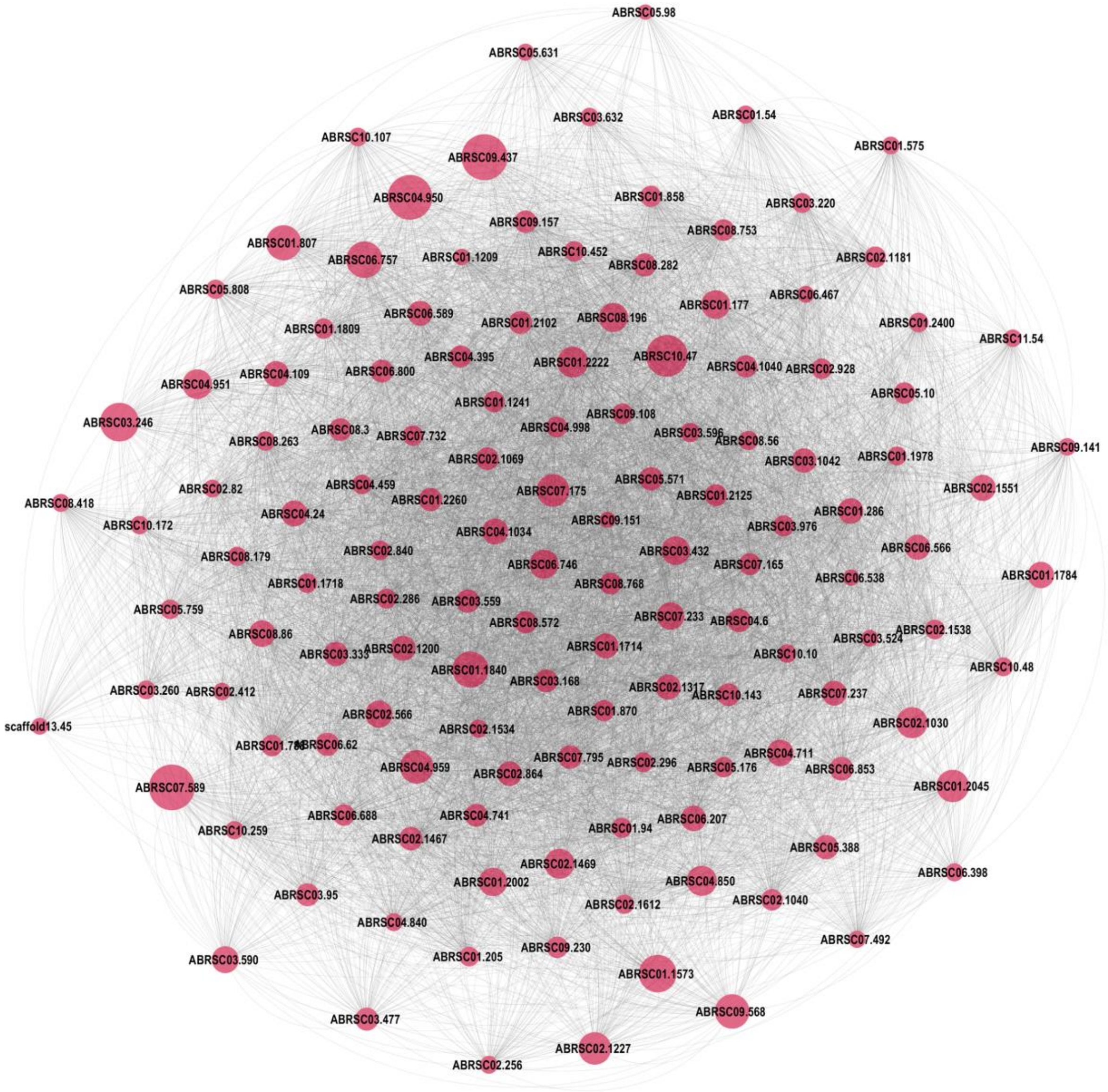


Figure S3: Gene co-expression network of genes in the darkred module

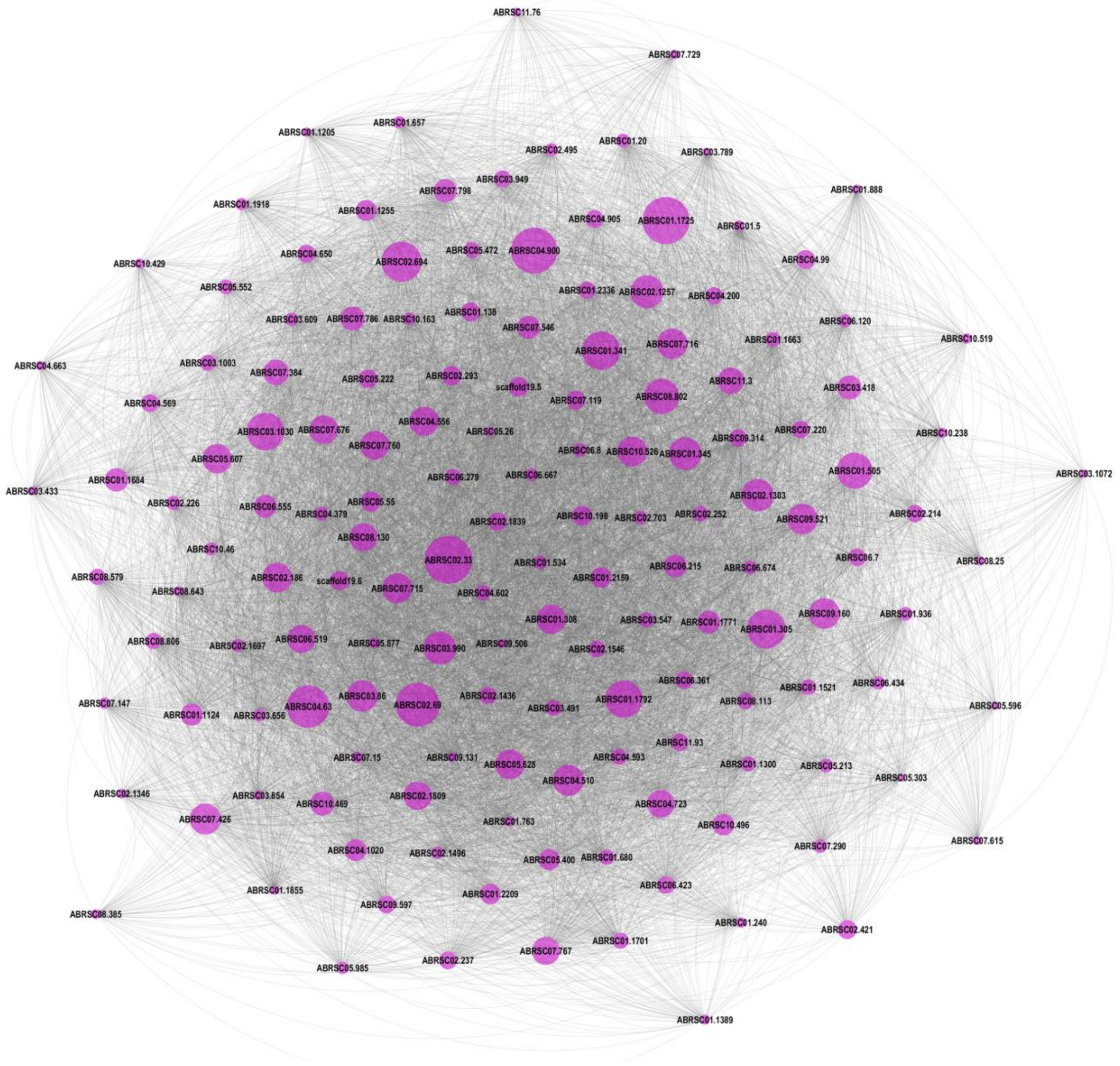


Figure S4: Gene co-expression network of genes in the magenta module

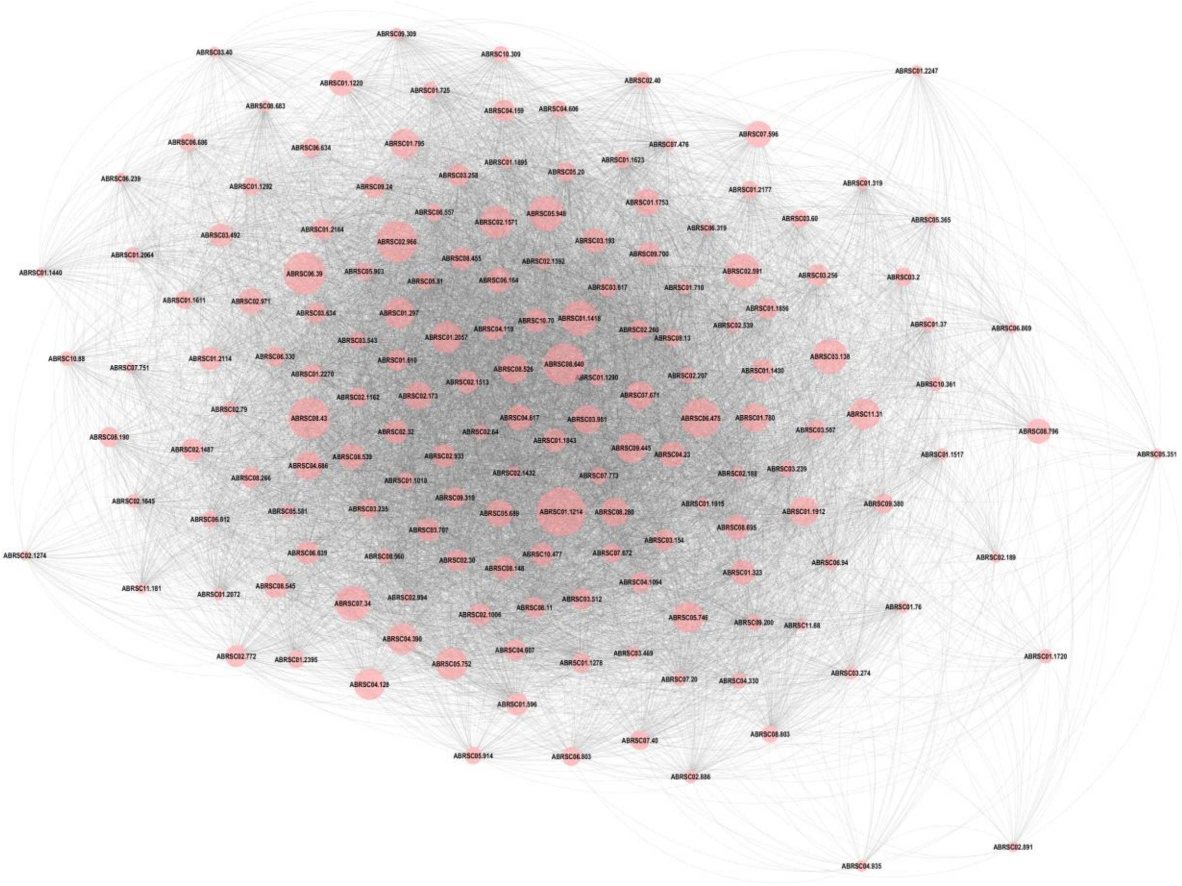

Figure S5: Gene co-expression network of genes in the salmon module

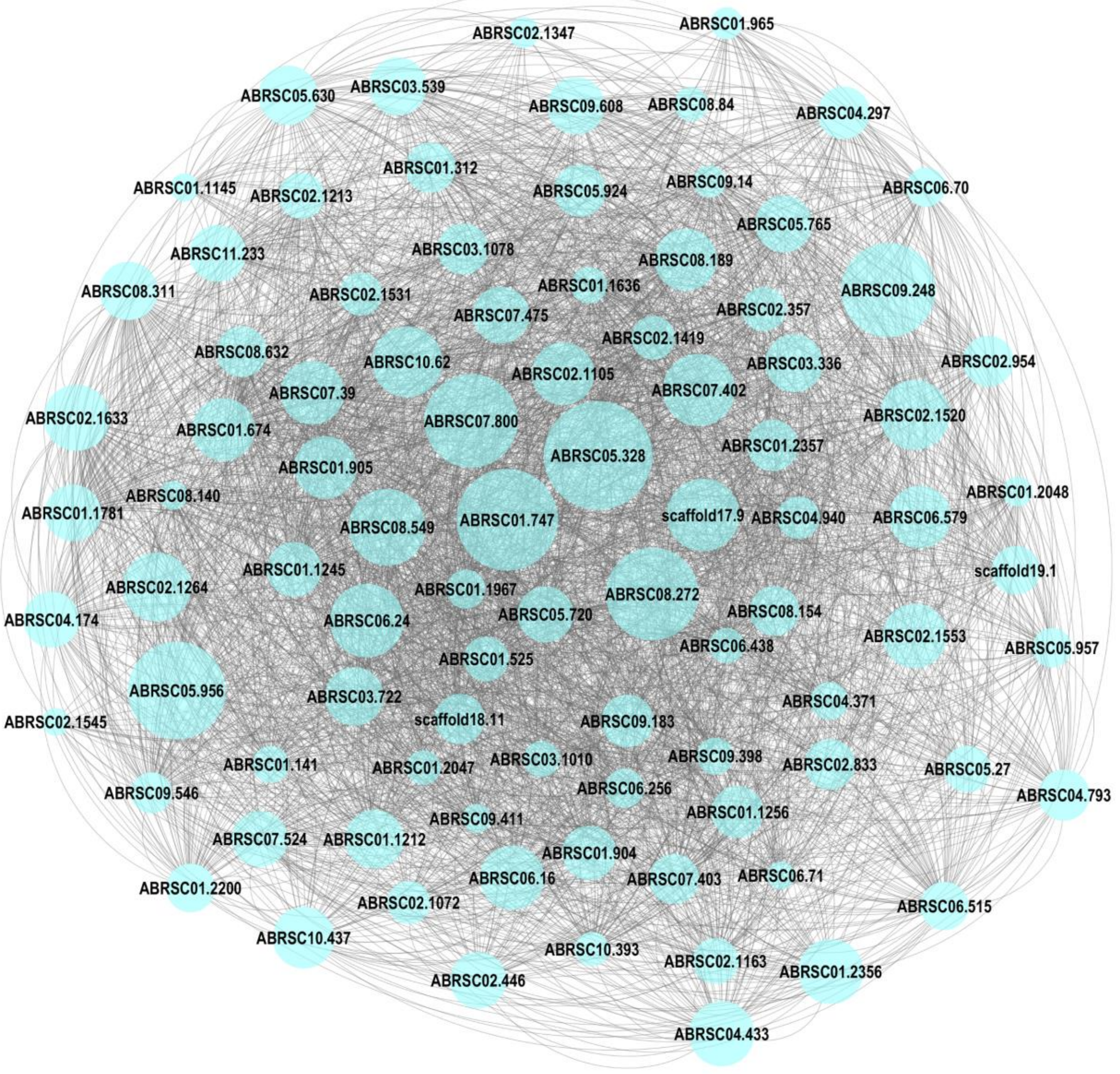


Figure S6: Gene co-expression network of genes in the lightcyan module

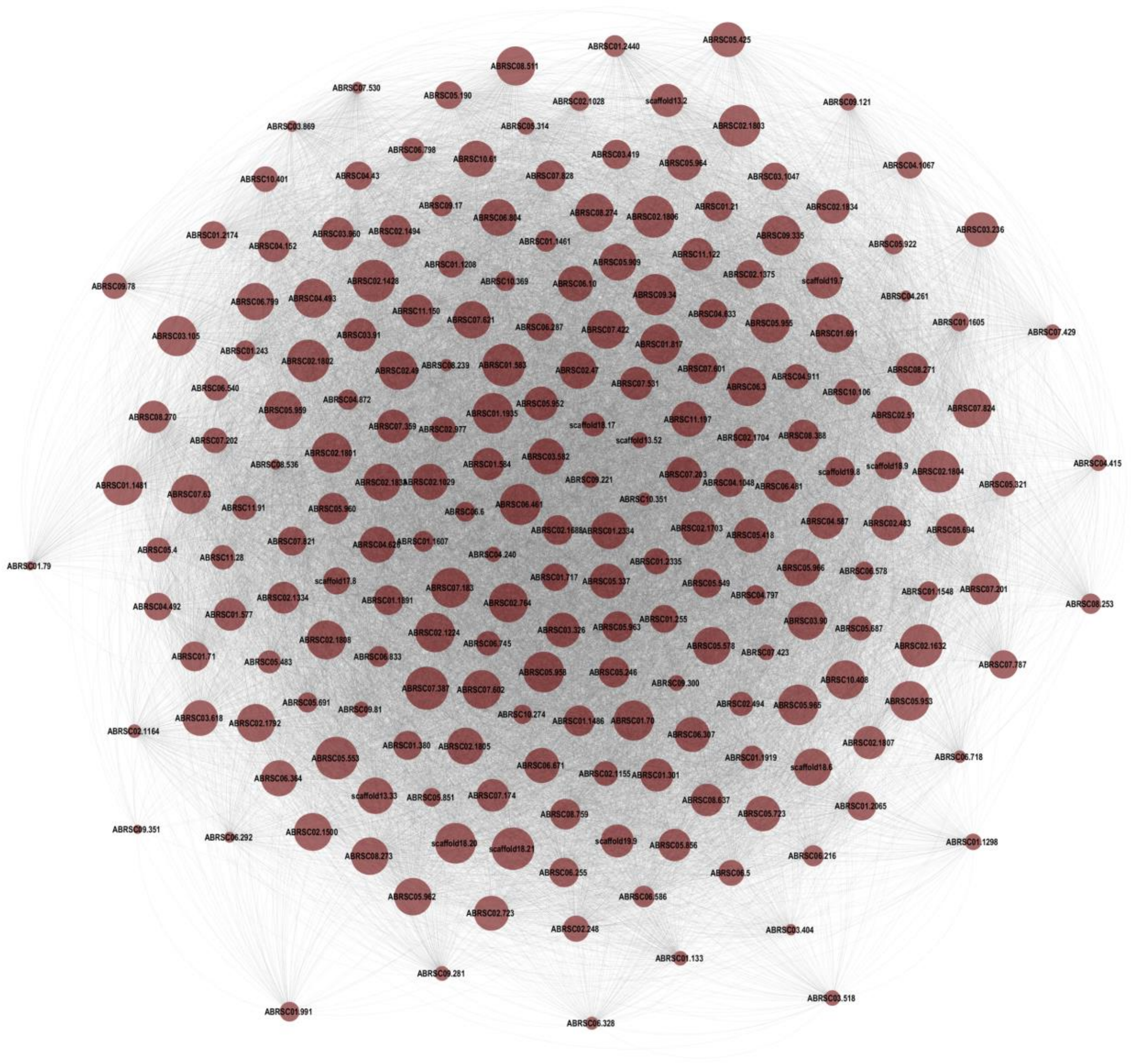

Figure S7: Gene co-expression network of genes in the brown module

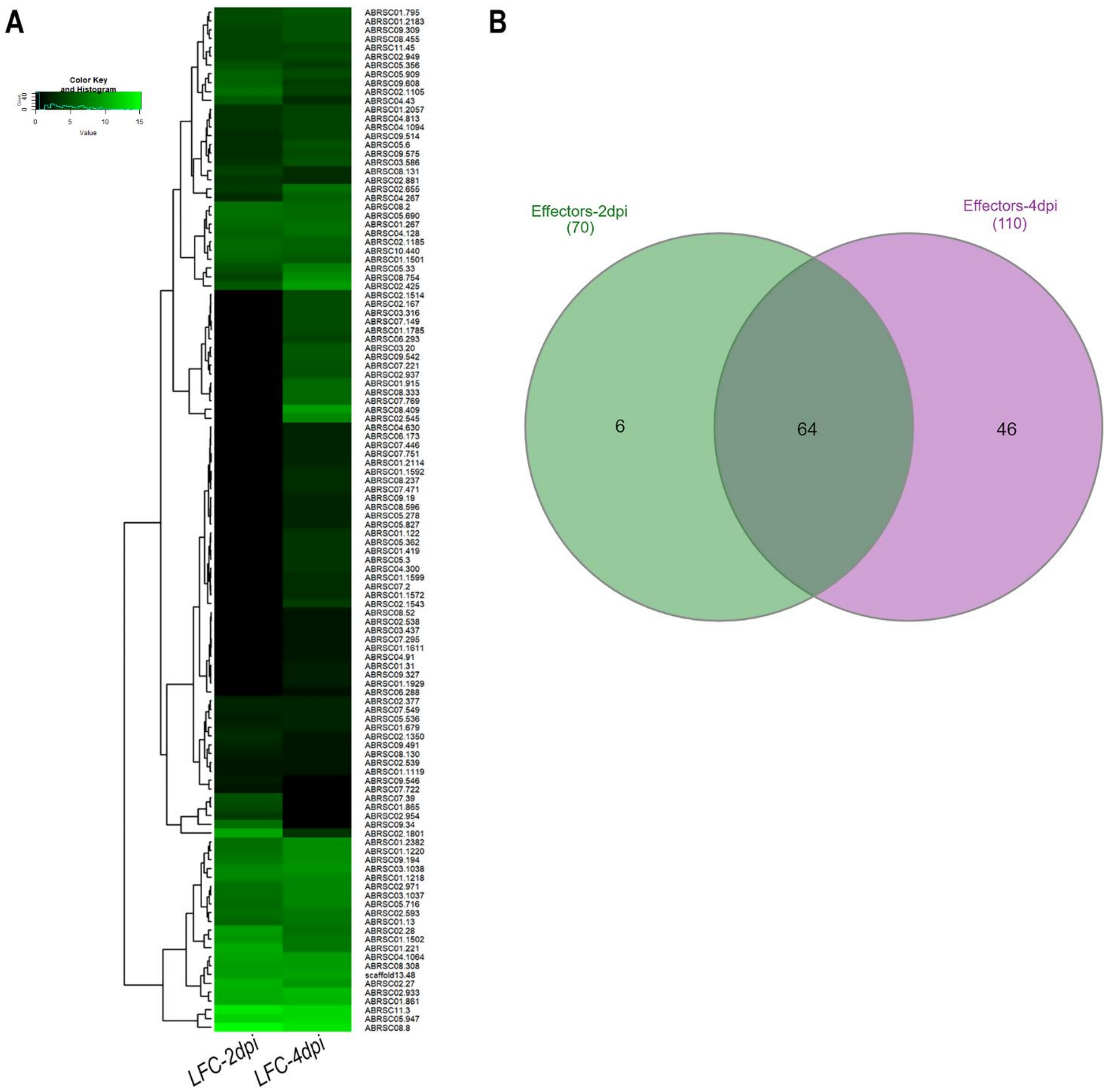


Figure S8: Gene expression profile of effectors of *A. brassicae*. (A) Heatmap depicting the $log_2$ FoldChange values of 116 effectors at 2 and 4 dpi. (B) Venn diagram representing the overlap between the effectors significantly upregulated at 2 and 4 dpi.
